# Integrative Transfer Network: Deep Transfer Learning Across Populations and Prediction Targets

**DOI:** 10.64898/2026.06.12.731936

**Authors:** Yan Gao, Yan Cui

## Abstract

Large-scale clinical and biomedical datasets increasingly contain both diverse subgroup attributes (e.g., demographic or clinical subgroups) and multiple prediction targets. Although various machine learning approaches can address subgroup differences or multi-target prediction, they often consider these aspects independently rather than jointly. To more effectively capture the shared and subgroup-specific information in such complex datasets, we propose the Integrative Transfer Network (ITN), a deep neural network designed to leverage data across subgroups and multiple related outcomes simultaneously. In extensive experiments, including time-to-event and classification tasks where demographic subgroups and multiple disease end-points are prevalent, ITN demonstrates consistent improvements in subgroup-specific prediction by borrowing strength from other subgroups and outcomes. We envision ITN as a unified framework for learning from heterogeneous datasets where subgroup-specific insights are critical.

## I. Introduction

The advent of large-scale, multi-faceted biomedical datasets has great potential to advance our understanding of complex diseases, necessitating the development of sophisticated analytical frameworks to decipher the underlying patterns and relationships effectively. These datasets are often characterized by interrelated outcomes and diverse subgroup populations, presenting a significant challenge for machine learning.

The emergence of integrative transfer networks (ITNs) heralds a significant advance in addressing these challenges. ITNs leverage deep learning architectures to enable transfer learning across different subgroups and multiple related outcomes, facilitating a more nuanced and comprehensive analysis. By integrating data across diverse demographic segments and related disease metrics, ITNs aim to enhance prediction accuracy for subgroup-specific outcomes, thereby addressing the critical issue of data distribution discrepancies that often impair the performance of traditional models.

This innovation is particularly crucial in the context of precision medicine, where understanding the subtle variations within and across patient subgroups can lead to more tailored and effective treatments. The application of ITNs across various clinical scenarios, ranging from cancer prognosis to cardiovascular disease outcomes, underscores their versatility and potential to transform our approach to disease prediction and management.

By addressing the limitations of existing methodologies and harnessing the full potential of complex biomedical datasets, integrative transfer networks represent a pivotal step forward in our quest for more personalized and effective healthcare solutions. Their development reflects a growing recognition of the intricate interplay between genetic, environmental, and lifestyle factors in disease progression and highlights the need for analytical frameworks that can navigate this complexity with sophistication and precision.

Biomedical datasets with multiple related outcomes and subgroup compositions have become increasingly common. Multiple related outcomes is a scenario where each sample in the dataset has different compatible and relevant labels, including comprise multi-faceted endpoints and comorbidity in disease study. Multi-faceted endpoints provide a measurement of disease outcomes from different perspectives, like the four different endpoints in TCGA cohort [1]. Comorbidity studies the coexistence of multiple diseases or disorders in relation to a primary disease in a patient [2], like different cardiovascular disease (CVD) related outcomes in the Sleep Heart Health Study (SHHS) [3], [4]. Subgroup composition is also common, in which each group which characterizing an attribute of each patient including age, sex, ethnics ancestry, education category or other factors.

Subgroup-specific prediction is important because of the distribution discrepancy between different groups have been widely observed [5]–[10]. This data distribution discrepancy may jeopardize machine learning performance such that the model trained on the whole population can not guarantee well optimized for each subgroup. Rolf etc. conducted a comprehensive study with data from multiple subgroups and highlighted the importance of subgroup allocations in training data [11]. Gao and Cui showed that the current mixture and independent learning schemes could result in unbalanced performance for different ethnic groups [12], [13]. Subgroup-specific model may lead to improved prediction performance on each subgroup and also be useful for reducing model performance disparities among demographic groups. Build a machine learning method for subgroup-specific prediction from dataset with multiple related outcomes and subgroup composition is challenge. Previous works handle these two issues separately. Subgroup composition is handled with independent learning or transfer learning, and multiple related outcomes are handled with multi-task learning [14] or regarded as competing risk [15]–[18], where each outcome is mutually exclusive. Therefore, there is an information loss in these models, not fully utilize the information in the datasets.

In this paper, we proposed a method, Integrative Transfer Network (ITN), which can be use for subgroup-specific prediction in datasets with subgroup composition and multiple related outcomes. For a specific outcome prediction of a certain subgroup, data from another subgroup and data from related outcomes (auxiliary outcomes) will be used to train a model jointly. During the testing, only the input feature of a certain subgroup is needed, and the auxiliary outcomes are not necessary for model evaluation. Using a comprehensive study of experiments with sex and genetic ancestry compositions, we show that ITN could achieve an improved performance in sex and racial-specific prediction. We also show that ITN outperforms three basic learning schemes in machine learning tasks of time-to-event prediction and classification. We anticipate ITN as a starting point for investigating methods for dataset with subgroup composition and multiple related outcomes.

## II. Related Work

### Deep Transfer Learning

Deep transfer learning can be classified into four categories: Instances-based, mapping-based, Network-based, and Adversarial-based [19]. *Instance-based* deep transfer learning methods use adjusted weights for the source domain and select a portion of instances that are to be added to the training set of the target domain with appropriate weights for knowledge transfer. [20] developed a transfer learning method (Festra) for inter-regional sandstone microscopic image classification by selecting significant divergence features between the regions and using an enhanced TrAdaBoost for instance transfer. [21] proposed a metric transfer learning framework (MTLF) with adjusted weight for instances to bridge the distributions discrepancy between different domains and used Mahalanobis distance to maximize the intra-class distances and minimize the inter-class distances for the target domain. The intention of *Mapping-based* deep learning models is to map instances from the source and target domains into an embedded space using domain alignment and adaptation methods. [22] proposed a classification and contrastive semantic alignment (CCSA) transfer learning method for image classification by maximizing the separation loss and minimizing the domain adaptation loss. [23] developed a convolutional neural network (CNN)-based model for transfer learning by using an adaption layer and an additional domain confusion loss to learn domain invariant representations. *Network-based* transfer learning methods refer to models that reuse a partial network trained from the source domain as a sub-structure for the target domain model. [24] designed a model for visual recognition by reusing some layers from the pretrained ImageNet to get mid-level representation for target domain images. [25] proposed a transfer learning to learn transferable features and jointly learn adaptive classifiers from source domain (labeled data) and target domain (unlabeled data) with a shared classifier. *Adversarial-based* methods use the generative adversarial net (GAN) technology to find an embedded space where the representation of both source and target domains are well-aligned [26]. [27] used a discriminator gate with adjusted weights to filter samples from the source domain that could cause negative transfer. [28] proposed a model called adversarial discriminative domain adaptation (ADDA) that combines discriminative modeling, untied weight sharing, and a GAN loss for transfer learning.

### Deep Survival Analysis

Deep learning methods have been used for survival analysis and show improved performance when compared to traditional Cox models. [29] proposed a DeepSurv model for survival prediction and treatment recommendation. [30] and [31] proposed a deep learning-based model (Cox-nnet) to predict the likelihood of patient survival from the high-dimensional mRNA transcription feature in the TCGA cohort. [32] used a deep learning model for analysis of kidney graft survival. [33] proposed a deep learning-based method for survival analysis with competing risks. [34] proposed a meta-learning model to improve the survival prediction performance of rare diseases from mRNA transcriptome data from the TCGA cohort. [35] proposed a generative deep learning approach for survival analysis using electronic health record (EHR) data. [36] proposed a deep recurrent model (RNN-SURV) for patient level survival analysis and achieved a better concordance index than other approaches. [37] proposed a survival analysis model using deep learning and active learning with a sampling module to obtain accurate prediction from clinical features using labeled and unlabeled data and evaluated their model using prostate cancer data from SEER-Medicare. [38] developed a deep convolutional neural network (DeepConvSurv) for survival analysis using pathological images from the screening trial lung cancer data. [39] proposed a deep learning-based model with global average pooling and a locality-constrained linear coding, Cox proportional hazards module, and a biomarker interpretation module for survival analysis, [40] developed an unsupervised deep learning-based residual convolutional auto encoder for survival analysis using unlabeled image data from patients. [41] proposed a deep learning-based feature integration methods for breast cancer survival prediction using multi-omics data with a concatenation autoencoder to preserve modality-specific information and a cross-modality autoen-coder to achieve modality-invariant representations.

## III. METHOD

### A. Problem Definition

In this section, we describe our ITN model for time-to-event prediction, and the model used for classification will have a similar setting. To begin with, we have a training set with multiple targets 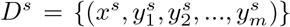 from the source domain, and we formulate it as 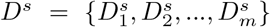, where 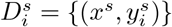 is the *i*^*th*^ prediction task in the source domain. For each 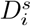, its feature *x*^*s*^ is drawn from a random distribution *X*^*s*^, and 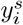 denotes the *i*^*th*^ clinical outcome for *x*^*s*^. In addition, we have a training set *D*^*t*^ from the target domain with the same composition, 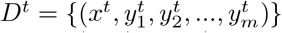, which can be formulated as 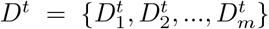, and 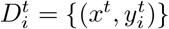 is the *i*^*th*^ prediction task in the target domain. Similarly, feature *x*^*t*^ is drawn from a random distribution *X*^*t*^, and 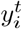 denotes the *i*^*th*^ clinical outcome of the target group. For each task in both domains, the *i*^*th*^ outcome *y*_*i*_ is a realization of a random variable *Y* (*y*_*i*_ *ϵ Y* ): for classification, *Y* is a discrete space with a certain number of categories; for regression, *Y* is a continuous variable; and in time-to-event prediction, *Y* is drawn from a paired distribution *Y ∼* (*T, E*) where *T* is a continuous variable for the event time and *E* is a binary variable that refers to the event status. Thus, the input datasets can be formulated in Equation 1, where *X*^*s*^ is referred as the source domain, and *X*^*t*^ is the target domain. For the *i*^*th*^ prediction target, 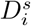 and 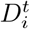 are the specific tasks in the source and target domains, 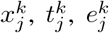 are the input feature, survival time, and event status of the *j*^*th*^ patient in group *k, M* and *N* are the number of samples in 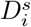 and 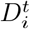. Under this setting, the objective of a specific time-to-event task 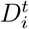 from the target domain is to find an optimal predictive function *f*_*i*_ : *f*_*i*_(*X*^*t*^) *→ T, E* using the knowledge from *D*^*s*^ *∪ D*^*t*^.

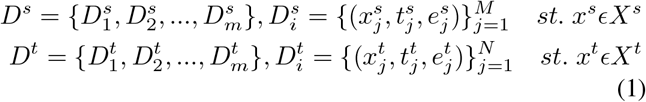

We assume that some tasks are compatible correlated and there is a distribution shift between *X*^*s*^ and *X*^*t*^. The assumption of relevance indicates that there exist a pair *h* and *i*, such that *D*_*h*_ and *D*_*i*_ share a common or close conditional distribution in both domains: 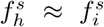 and 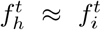. In addition, the domain shift assumption suggests that there is distribution difference across domains, *P* (*X*^*s*^) ≠ *P* (*X*^*t*^). The problem formulated here is a mixture of multi-target and transfer learning problem, which contains a supervised domain adaptation and a multi-target learning component. In this work, we are specially focused on the scenarios where the target task has related tasks available in both domains. We aim to work on scenarios where each data point has multiple targets in both domains and the total number of samples in each domain are limited. Such scenarios are much common in multi-faced disease and comorbidity.

### B. Baseline Deep Cox Neural Network

In our paper, we used a deep Cox neural network [42] as the baseline prediction model for each given task *D*_*i*_ from both domains. Here, we initiate *D*_*i*_ as *D* = {(*x, T, E*)}, where *x* is the input feature, *T* denotes the event time, and *E* refers to the event status of a group of patients. The baseline model is a mixture of deep learning structure followed with a Cox regression layer. The deep learning structure is used to map the high-dimensional input feature *X* into an embedding space *Z*, and the Cox regression layer is used to make a prediction from *Z*. To avoid overfitting, we used dropout layers, learning rate decay, and a *L*_2_ regularization term.

### C. Integrative Transfer Network (ITN)

For a given dataset with two subgroups and two related TTE outcomes which can be formulated as Equation 2, where 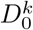 and 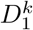 are the main task and auxiliary task for domain *k, x*^*s*^ and *x*^*t*^ are the input features from both domains, 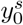 and 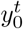 are the main predicting target and 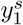 and 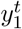 are the auxiliary predicting target. Here, we focus on the main prediction task in the target domain, 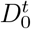, and model other tasks with the same manner. Therefore, the objective of ITN is to to learn a predictive function using all information from *D*.

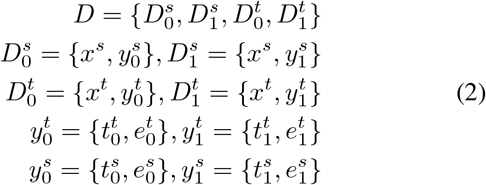

The architecture of ITN is shown in Fig III-A. In the Figure, *F* is the feature extractor whose purpose is to align the distribution of each task feature in the embedded space, *z* and *z*^*′*^ are the embedding of the target and the source domains, *C*^*t*^ is the Cox regression module for the main target in the target domain, *y*^*′*^ is the output risk of *C*^*t*^, *C*^*s,t*^ is the Cox module for the auxiliary target in both domains, 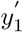 is the output of 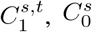, is the Cox module for the main target in the source domain and 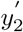 is the output of 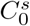, *D* is a discriminator, 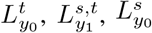, are the loss of three TTE tasks: the main task in the target domain, the auxiliary task for both domains, and the main task in the source domain, *L*_*ADV*_ is the adversarial loss of two domains, *λ*_0_, *λ*_1_, *λ*_2_, *λ*_3_ are the adjusted weights. To reduce parameters in our model, we set *λ*_1_ = *λ*_2_ = *λ*_3_ = 1 *−λ*_0_.

The overall objective function of ITN *L*_*IT N*_ is a linear combination of *L*_*MT*_ and a domain adversarial loss *L*_*ADV*_, as shown in Equation 3. *L*_*MT*_ is a weighted sum of three TTE loss terms, 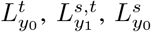 as shown in Equation 4. 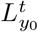 is a regularized partial hazard loss of 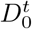 as shown in Equation ref 5, where 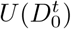 is the set of uncensored samples in 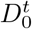,*β*_1_ is the weight of 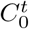, *W*_1_ is the weight of *F* and 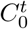, and *λ*_1_ is the weight of the *L*_2_ regularization term. Similarly, 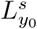 and 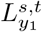 are the partial hazard loss of 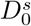 and 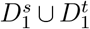, as shown in Equation 6 and 7. *β*_2_ is the weight of 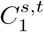, *W*_2_ is the weight of *F* and 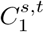, *β*_3_ is the weight of 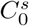, *W*_3_ is the weight of *F* and 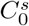. The domain loss *L*_*ADV*_ is the binary cross-entropy loss as defined in Equation 8, where *F* is the feature extractor and *D* is the discriminator as shown in Fig III-A, *x*_*s*_ and *x*_*t*_ are the input features from the source and the target domains.

**Fig. 1.**
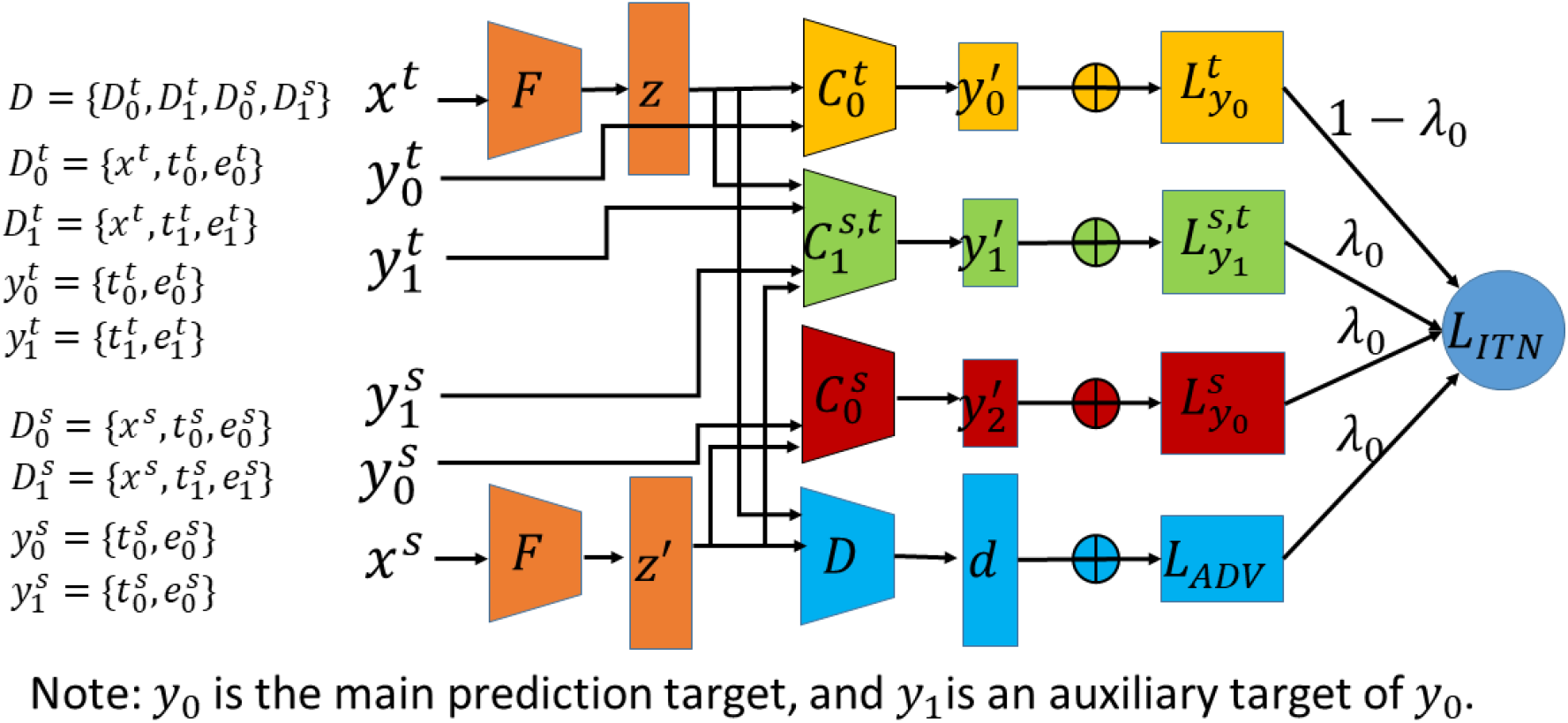
The architecture of ITN. During the training stage, the loss of the main target in the target domain, the related target from both domains, and main target from the source domain were optimized jointly, the GAN loss maximizes the distances between two domains.

The prediction function of the 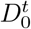 can be decomposed into 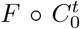. Similarly, the prediction function of the auxiliary target 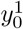 and 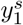 can be decomposed into 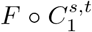, and the prediction function of 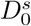 can be decomposed into 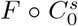. The discriminator *D* is used to handle the distribution shift between *x*^*s*^ and *x*^*t*^, to ensure that the feature distribution of *P* (*z*) and *P* (*z*^*′*^) can not be distinguished by in the embedding space. Under this setting, the ITN would learn knowledge from related targets jointly and transfer knolwdge from the source domain.

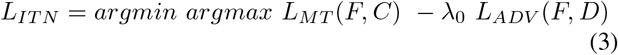

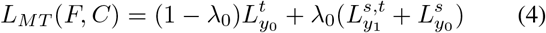

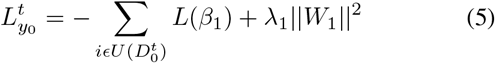

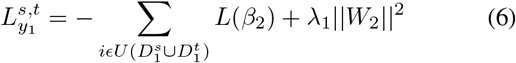

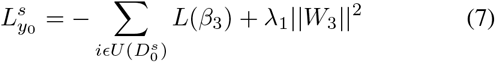

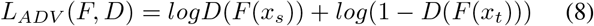

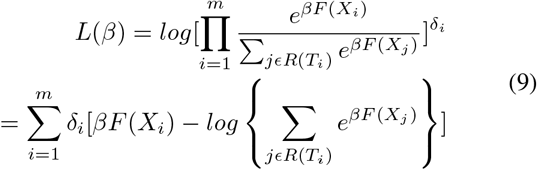

### D. Hyperparameter Tuning

The training time for our ITN model requires an extensive hyper-parameter searching, including deep neural network related parameters and our defined parameters. We employed a Bayesian optimization to identify the parameters for network structure, including the number of layers, the number of nodes in each layer, learning rate, momentum, dropout rate, and activation functions. During the optimization, we reserved 20% of the data from the training set as the validation set, and trained the desired model weights using the hyper-parameters suggested by Bayesian optimization. After the optimization process, the best network structure will be used to retrain a deep learning model.

## IV. Experiments

We conducted experiments with machine learning tasks assembled from three datasets: 1) The Cancer Genome Atlas (TCGA) ^1^, 2) Acute Kidney Injury (AKI) Study [43], 3) Sleep Heart Health Study (SHHS) [3], [4]. All of these datasets are public available and do not contain any personal identifiers or offensive contents. In our experiments, we focused on subgroups based on sex and genetic ancestry, which subgroup-specific disparities have been widely observed [5], [6], [44]–[46], [7]–[10].

### A. Experimental Setup

For each machine learning task, we compared the prediction performance of each subgroup from our ITN with three baseline machine learning schemes: mixture learning, independent learning, and naive transfer. The training set for three baseline learning schemes is different: mixture learning uses the set of 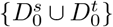, independent learning uses 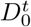, and naive transfer uses 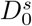. For ITN, the training set is 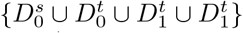. After model training, the testing set of 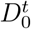 will be used for model evaluation.

For each learning scheme, we applied a 10-fold stratified cross-validation for training and testing split, in which samples are stratified on the basis of group label and event attributes. We performed 10 independent runs for each experiment with different random seeds and partitions. We evaluated each learning scheme for each experiment using the concordance index [47] and Area under the ROC curve (AUC).

### B. Experiment 1: Sex-specific Prediction of Cancer Outcomes

We studied the sex-specific cancer prognosis using the TCGA cohort, which contains omics data from more than 11,000 patients. We divided the cohort into 40 independent datasets by disease as shown in Appendix A. For each dataset, the input feature is the mRNA expression, and the prediction target are four different clinical endpoints: overall survival (OS), progression-free interval (PFI), disease-free interval (DFI), and disease-specific survival (DSS). As a result, we assembled 160 tasks from a combination of 40 cancer types and 4 clinical endpoints. In our experiments, OS and DSS are survival related tasks, PFI and DFI are progression related tasks. We applied four learning schemes to each sex group, as shown in Section IV-A. To study the male group, we set the female group as the source domain, and vice versa.

To handle missing values, we filtered these features with missing values for any disease to generate a total of 17,176 transcripts. These features are normalized to improve machine learning practice. To retain a meaningful study, we filtered the tasks with less than 50 samples in a sex group. We also removed those tasks have less than 10 observed events in a sex group. For each sex group, we filtered out the machine learning tasks that have a C-index less than 0.6, resulting a total of 29 tasks from 9 cancers and 4 different clinical outcome endpoints.

The sex-specific prediction for the male and female are shown in Table I and Table II. The left four columns show the basic task information, and the last four columns show the testing C-index from four different learning schemes. The performance varies from different learning schemes on both sex groups. For the male group, the ITN outperformed all other three learning schemes in 13 out of 15 tasks. For the female group, 10 out of 14 tasks showed a performance improvement using ITN. Both tables show clear potential of ITN in improving the performance for sex-specific prediction.

**TABLE I.** The c-index for cancer outcome prediction using different learning schemes (Male group)

| Cancer type | Endpoint | Events | Censored | Mixture | Independent | Naive | ITN |
| --- | --- | --- | --- | --- | --- | --- | --- |
| ACC | OS | 11 | 20 | 0.725 | 0.821 | 0.511 | **0.876** |
| GBMLGG | OS | 155 | 234 | 0.819 | 0.802 | 0.814 | **0.848** |
| GBMLGG | PFI | 189 | 200 | 0.766 | 0.747 | 0.738 | **0.784** |
| KIRC | DSS | 76 | 264 | **0.719** | 0.701 | 0.718 | 0.714 |
| ACC | PFI | 14 | 17 | 0.710 | 0.664 | 0.556 | **0.810** |
| KIRC | PFI | 115 | 229 | 0.666 | 0.654 | 0.654 | **0.689** |
| PAAD | DSS | 36 | 60 | 0.655 | 0.644 | **0.675** | 0.639 |
| KIRC | OS | 111 | 234 | 0.667 | 0.643 | 0.613 | **0.678** |
| UVM | PFI | 16 | 28 | 0.559 | 0.608 | 0.602 | **0.849** |
| PAAD | OS | 47 | 51 | 0.598 | 0.607 | 0.588 | **0.614** |
| BLCA | DSS | 87 | 205 | 0.598 | 0.605 | 0.504 | **0.660** |
| PAAD | PFI | 56 | 42 | 0.623 | 0.587 | 0.564 | **0.634** |
| BLCA | OS | 127 | 173 | 0.579 | 0.550 | 0.537 | **0.609** |
| SARC | OS | 41 | 77 | 0.618 | 0.549 | 0.600 | **0.702** |
| SARC | PFI | 64 | 54 | 0.596 | 0.548 | 0.574 | **0.644** |

**TABLE II.** The c-index for cancer outcome prediction using different learning schemes (Female group)

| Cancer type | Endpoint | Events | Censored | Mixture | Independent | Naive | ITN |
| --- | --- | --- | --- | --- | --- | --- | --- |
| GBMLGG | OS | 187 | 98 | 0.816 | 0.765 | 0.804 | **0.831** |
| ACC | OS | 31 | 17 | 0.692 | 0.410 | 0.570 | **0.817** |
| ACC | PFI | 21 | 27 | 0.703 | 0.562 | 0.592 | **0.789** |
| KIRC | DSS | 149 | 33 | **0.780** | 0.751 | 0.778 | 0.757 |
| GBMLGG | PFI | 155 | 130 | 0.734 | 0.696 | 0.722 | **0.741** |
| KIRC | PFI | 143 | 44 | 0.739 | 0.698 | **0.741** | 0.730 |
| KIRC | OS | 124 | 64 | **0.710** | 0.694 | 0.704 | 0.708 |
| UVM | PFI | 22 | 13 | 0.562 | 0.442 | **0.689** | 0.683 |
| SARC | OS | 84 | 57 | 0.653 | 0.559 | 0.526 | **0.680** |
| BLCA | OS | 56 | 51 | 0.625 | 0.567 | 0.586 | **0.662** |
| BLCA | DSS | 67 | 34 | 0.629 | 0.518 | 0.628 | **0.645** |
| BLCA | PFI | 59 | 48 | 0.585 | 0.562 | 0.553 | **0.634** |
| PAAD | PFI | 32 | 48 | 0.590 | 0.491 | 0.596 | **0.632** |
| SARC | PFI | 67 | 74 | 0.597 | 0.530 | 0.538 | **0.607** |

**TABLE III.**
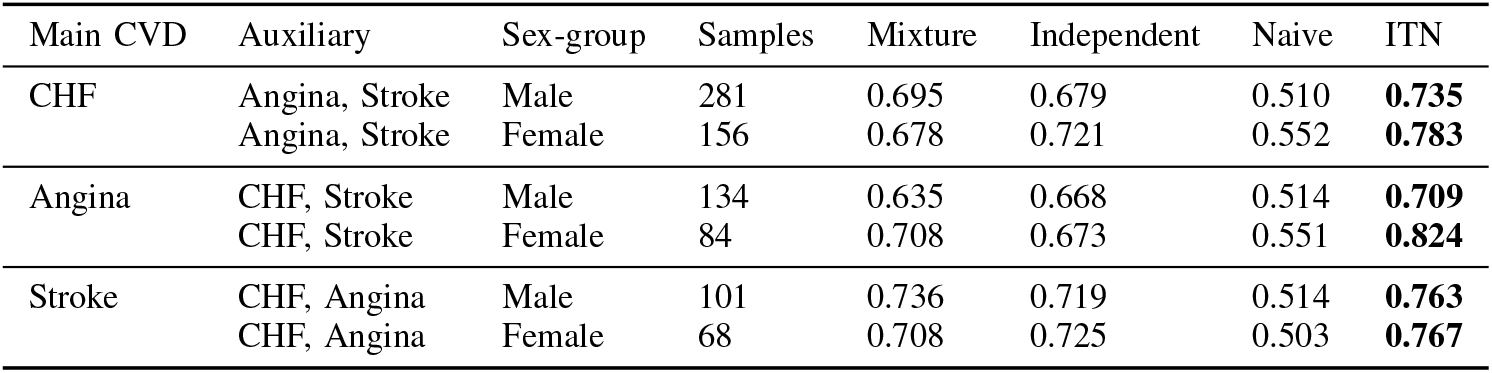
The c-index for time-to-event prediction of CVD outcomes using different learning schemes.

### C. Experiment 2: Sex-specific Prediction of Cardiovascular Disease Outcomes

We conducted a sex-specific prediction of cardiovascular disease (CVD) outcomes using the data from national Sleep Heart Health Study^2^. The dataset contains 1279 clinical features, and the follow-up time of 10 different CVD outcomes from 5804 records. We removed the records with 20% missing features and dropped the feature with missing values. We converted the *rcrdtime* (total recording time) into number of minutes. We filtered these samples with missing values in sex or event time. We further removed these diseases have less than 50 samples in a sex group, resulting three heart diseases: Angina (type 1) with 218 samples, congestive heart failure (type 4) with 437 samples, and Stroke (type 10) with 169 samples.

The prediction performance of each sex-group from different learning schemes are shown in Table III. Data from the table show that ITN outperformed all other learning schemes for both sex-groups in three events prediction. The best improvement occurred to the female group in Angina event, from 0.71 to 0.82.

### D. Experiment 3: Sex-specific Prediction of Acute Kidney Injury

We further adapted our method to a classification problem [43], to the Acute Kidney Injury (AKI) from electronic medical records. In the dataste, each patient has three binary case/control labels. Label 1 is defined as: *Can we predict AKI before its onset using data before the onset time?* Label 2 is defined as: *Can we predict at admission if AKI will occur for patients during their stay?* Label 3 is defined as: *Can we predict at admission if AKI will occur within various numbers of days afterwards?*

The AKI dataset contains 1917 features from clinical inquiries and laboratory tests. We filtered features with *≥*5% missing values and removed patients have missing data, resulting 208 features with 508 patients (108 cases) in perspective 1,300 patients (44 cases) in perspective 2, and 300 patients (10 cases) in perspective 3. We normalized each feature to best fit the deep learning practice. We expanded the race category into 4 features.

We focused on perspective 1 which has adequate case samples and data from label 2 and 3 is used as auxiliary tasks. We replaced the Cox modules with classification modules in ITN and changed the equation 9 to binary cross entropy to handle the classification problem. The AUCs for each learning scheme are shown in Table IV, and we noticed our ITN outperformed other learning schemes.

**TABLE IV.** The AUC for acute kidney injury prediction using different learning schemes.

| Sex-group | Case | Control | Mixture | Independent | Naive | ITN |
| --- | --- | --- | --- | --- | --- | --- |
| Male | 59 | 237 | 0.672 | 0.699 | 0.602 | **0.720** |
| Female | 49 | 163 | 0.654 | 0.680 | 0.592 | **0.702** |

### E. Experiment 4: Ancestry-specific Prediction of Disease Specific Survival Outcome

We also studied the ancestry-specific prediction in cancer prognosis using the uterine corpus endometrial carcinoma (UCEC) dataset from TCGA. The dataset contains 17176 mRNA expression features from 482 patients from the European American (EA) and African American (AA) racial groups, as well as the outcome of disease-specific survival (DSS). We used the overall survival of UCEC as the auxiliary task. The genetic ancestry of each patient is from a feature matching process [48]. The testing C-index of each racial group in UCEC DSS prediction is shown in Table V. From the table, we noticed ITN achieved the best performance and a significant improvement on the AA group. Compared with mixture learning, ITN slightly improved the performance of EA group, due to the limited samples size in the source domain (AA).

**TABLE V.** The c-index for ucec disease-specific survival outcome prediction using different learning schemes (EA and AA groups)

| Ancestry | Censorship | Events | Mixture | Independent | Naive | Ours |
| --- | --- | --- | --- | --- | --- | --- |
| EA | 336 | 36 | 0.715 | 0.676 | 0.548 | **0.723** |
| AA | 96 | 16 | 0.623 | 0.403 | 0.605 | **0.681** |

**TABLE VI.** The abbreviations of tcga cancer types.

| Abbreviation | Cancer Type |
| --- | --- |
| ACC | Adrenocortical carcinoma |
| BLCA | Bladder urothelial carcinoma |
| BRCA | Breast invasive carcinoma |
| CESC | Cervical and endocervical cancers |
| CHOL | Cholangiocarcinoma |
| COAD | Colon adenocarcinoma |
| DLBC | Lymphoid Neoplasm Diffuse Large B-cell Lymphoma |
| ESCA | Esophageal carcinoma |
| GBM | Glioblastoma multiforme |
| HNSC | Head and Neck squamous cell carcinoma |
| KICH | Kidney Chromophobe |
| KIRC | Kidney renal clear cell carcinoma |
| KIRP | Kidney renal papillary cell carcinoma |
| LAML | Acute Myeloid Leukemia |
| LGG | Brain Lower Grade Glioma |
| LIHC | Liver hepatocellular carcinoma |
| LUAD | Lung adenocarcinoma |
| LUSC | Lung squamous cell carcinoma |
| MESO | Mesothelioma |
| OV | Ovarian serous cystadenocarcinoma |
| PAAD | Pancreatic adenocarcinoma |
| PCPG | Pheochromocytoma and Paraganglioma |
| PRAD | Prostate adenocarcinoma |
| READ | Rectum adenocarcinoma |
| SARC | Sarcoma |
| SKCM | Skin Cutaneous Melanoma |
| STAD | Stomach adenocarcinoma |
| TGCT | Testicular Germ Cell Tumors |
| THCA | Thyroid carcinoma |
| THYM | Thymoma |
| UCEC | Uterine Corpus Endometrial Carcinoma |
| UCS | Uterine Carcinosarcoma |
| UVM | Uveal Melanoma |
| COADREAD | Colorectal adenocarcinoma (COAD+READ) |
| GBMLGG | Glioma (GBM+LGG) |
| STES | Stomach and Esophageal carcinoma |
| PanGyn | Pan-gynecological cancers (OV+CESC+USC+UCEC) |
| PanGI | Pan-gastrointestinal cancers (LUSC+HNSC+ESCA+CESC+BLCA) |
| PanSCCs | Pan-squamous cancers (COAD+READ+STAD+ESCA) |
| KIPAN | Pan-kidney cancers (KICH+KIRC+KIRP) |

## V. Conclusion and Discussion

In this work, we developed ITN, an integrative transfer learning model for subgroup specific prediction. The method has shown significant improvement in time to event prediction and classification. ITN outperforms three baseline machine learning schemes by incorporating data from multiple modalities, including related outcomes and a similar domain. Such method would be useful for dataset with subgroups and multiple compatible related outcomes. The method is also useful for addressing the important challenges such as reducing the negative impacts of data inequality among different demographic groups on machine learning performance, and thus help to improve social equity. We expect that the method has broad applications in disease prognosis and beyond.

## VI. Future Works

In this paper, we study the performance of sex and racial specific survival analysis using different machine learning schemes. We show that our method would achieve an improved performance for both sex-based and racial-based groups. In the future, we will study more diseases, incorporate more features, and investigate the model explainbility and model fairness on each subgroup.

## Acknowledgments

This work was supported by the US National Cancer Institute grant R01CA262296.

## Footnotes

1 The Cancer Genome Atlas Program. (https://www.cancer.gov/aboutnci/organization/ccg/research/structural-genomics/tcga)

2 Sleep Heart Health Study, https://www.sleepdata.org/

## Notes

### Competing Interest Statement

The authors have declared no competing interest.

